# TerC Proteins Function During Protein Secretion to Metalate Exoenzymes

**DOI:** 10.1101/2023.04.10.536223

**Authors:** Bixi He, Ankita J. Sachla, John D. Helmann

## Abstract

Cytosolic metalloenzymes acquire metals from buffered intracellular pools. How exported metalloenzymes are appropriately metalated is less clear. We provide evidence that TerC family proteins function in metalation of enzymes during export through the general secretion (Sec-dependent) pathway. *Bacillus subtilis* strains lacking MeeF(YceF) and MeeY(YkoY) have a reduced capacity for protein export and a greatly reduced level of manganese (Mn) in the secreted proteome. MeeF and MeeY copurify with proteins of the general secretory pathway, and in their absence the FtsH membrane protease is essential for viability. MeeF and MeeY are also required for efficient function of the Mn^2+^-dependent lipoteichoic acid synthase (LtaS), a membrane-localized enzyme with an extracytoplasmic active site. Thus, MeeF and MeeY, representative of the widely conserved TerC family of membrane transporters, function in the co-translocational metalation of Mn^2+^-dependent membrane and extracellular enzymes.

## Introduction

Metal ions are essential for life, in large part due to their roles as cofactors for enzymes where they can serve as an electrophilic center or redox catalyst^1, 2^. Metalloenzymes most often function with a specific metal that is acquired during protein folding or by binding of metal to an already folded apo-protein. Some metals bind proteins with high affinity and exchange slowly if at all. Others bind more loosely and exchange frequently. These properties have been summarized in the Irving-Williams series: Mn(II)<Fe(II)<Co(II)<Ni(II)<Cu(II)>Zn(II)^3^. The most abundant metals in the cytosol (Mn, Fe, and Zn) are generally present as divalent ions and are referred to here without reference to ionic state. Cytosolic enzymes acquire metals from a buffered pool^4, 5^, with Mn and Fe at low micromolar levels^6, 7^. For high affinity metals, such as Cu and Zn, enzymes may require a metallochaperone for metal insertion^8^.

Metalation of enzymes with active sites external to the cell membrane is not as well understood. Zn-requiring enzymes may acquire this ion from the environment^9, 10^. However, metalation of enzymes that require lower affinity metals is more problematic. For example, if a Mn-requiring enzyme is secreted from the cell without an associated metal ion it can easily be mismetallated by Cu or Zn^9^. One solution is to metalate the protein inside the cell and secrete the folded metalloprotein through the TAT-dependent secretion system^9^. However, this is not a general solution and many metalloproteins are exported in an unfolded state through the SecYEG-dependent general secretion pathway. How these exported proteins are properly metalated has not been resolved.

*Bacillus subtilis* is an important biotechnology platform often employed to produce secreted proteins^11, 12^. Most secreted proteins transit the membrane through the heterotrimeric SecYEG translocon driven by the SecA ATPase^13^. In *Escherichia coli*, a larger complex, the holotranslocon, comprises SecYEG together with SecDF-YajC and a YidC membrane protein insertase^14^. Further association with a variable set of folding chaperones and quality control proteases defines a larger secretosome complex^15^ that may also include the F_1_F_o_ ATPase^16^.

Here we report that the *Bacillus subtilis* TerC proteins MeeF(YceF) and MeeY(YkoY) are involved in <u>m</u>etalation of <u>e</u>xo<u>e</u>nzymes with Mn, an important cofactor for diverse enzymes^17, 18^. TerC proteins (Pfam03741) are poorly understood membrane proteins previously implicated in Mn export^19, 20^. Proteomic and genetic studies indicated that TerC proteins interact with the secretosome, suggestive of a role in co-translocational protein metalation. Mutants lacking these proteins (11*meeF* 11mee*Y*=FY mutants) were defective in protein secretion and in metalation of LtaS, a Mn-dependent enzyme that synthesizes membrane-associated lipoteichoic acids polymers. Consistently, the FtsH protease, critical for clearing jammed translocons in the membrane^21^, was essential in the FY strain, and overexpression of FtsH improved fitness of the FY mutants. Our results implicate TerC proteins as accessory subunits of the holotranslocon that mediate the metalation of exoenzymes. A similar biochemical role may explain phenotypes resulting from mutations in the related plant^22^, yeast^23^, and human^24^ orthologs.

## Results

### Cells lacking the two major TerC proteins (MeeF and MeeY) are defective in production of extracellular proteases

The roles of TerC proteins are enigmatic. Double mutants lacking both *meeF(yceF)* and *meeY(ykoY),* designated as FY mutants, display increased Mn accumulation under conditions of excess Mn, suggestive of a role in Mn export^20^. However, this role is secondary to the Mn exporters MneP and MneS cation diffusion facilitator proteins^20, 25^. Thus, the role of TerC proteins in Mn homeostasis is unclear.

The FY double mutant, but not the single mutants, displayed a large (74%) decrease in colony size on LB agar (Fig. 1a, Fig. S1a), suggesting that MeeF and MeeY (∼40% aa identity) have overlapping functions. In contrast, the doubling time of WT, F, Y and FY mutants was comparable in liquid LB medium (Fig. S1b). The Mn concentration in LB medium (<0.2 µM) is sufficient to support normal growth^25^, but far below toxic levels (200 μM)^20, 26^. Thus, the reduced fitness of the FY mutant is unlikely to be related to Mn detoxification.

**Figure 1.**
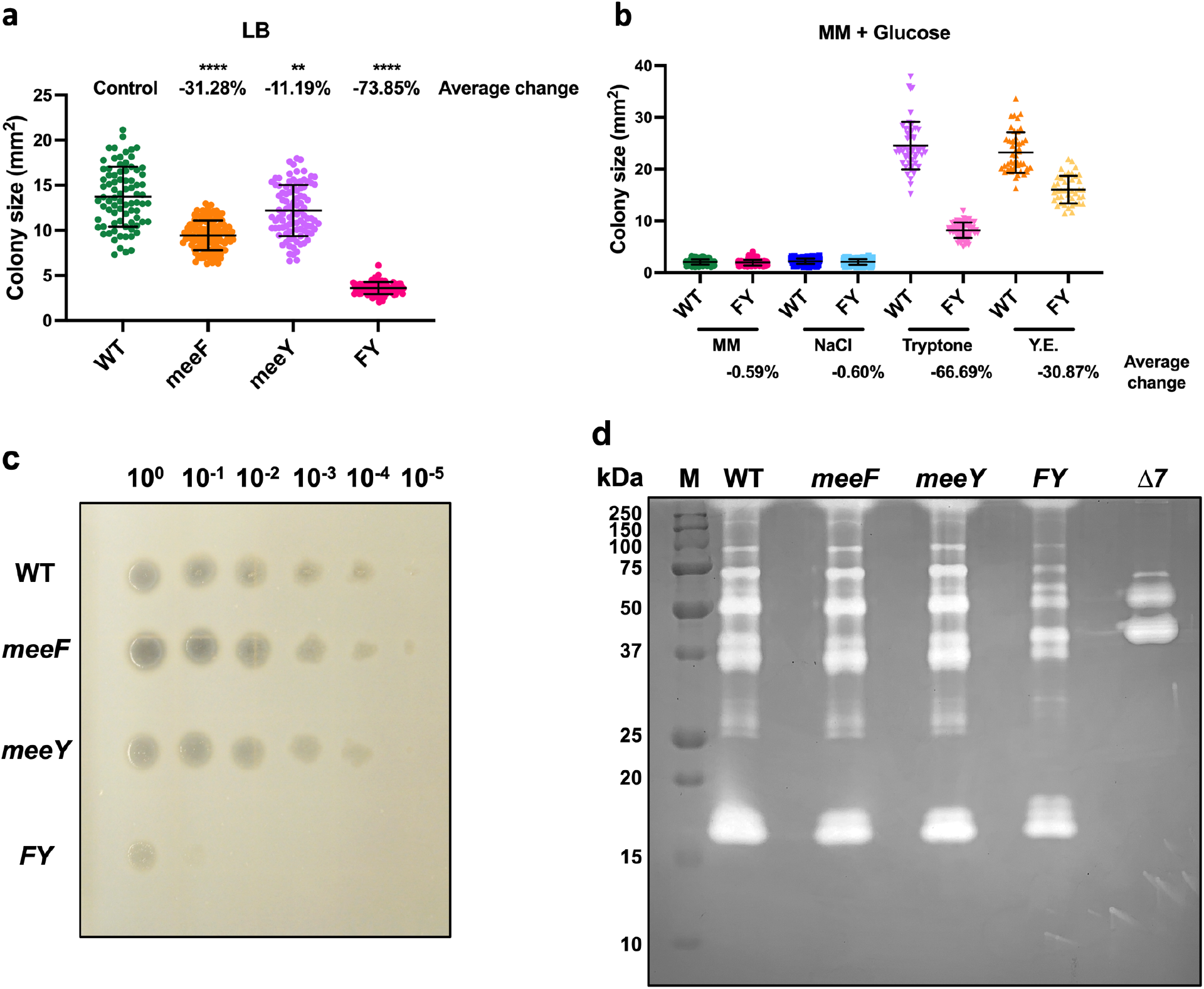
MeeF and MeeY are required for efficient secretion of feeding proteases to access nutrients in tryptone. (**a**) Colony size of WT, *meeF, meeY, and* FY on LB agar. (**b**) Colony size of WT and FY mutant on a defined glucose-minimal media (MM). MM agar plates were made with and without NaCl, tryptone, or yeast extract (Y.E.). Agar plates with well isolated colonies were imaged after 24 h at 37°C, and sizes measured using Image J. 40 or more isolated colonies from at least three independent cultures were included in the measurements for each strain, data is presented as mean ± SD. **, P=0.0014; ****, P<0.0001, P value was calculated using Welch’s t test, two-tailed. Average changes were calculated as “change = (sample – control) / control * 100%”. WT is the control for each nutrient component. (**c**) Protease hydrolysis of different strains on 5% milk agar plates. Zone of clearance due to protease activity was imaged. (**d**) Extracellular protease activities in the supernatants were detected by gelatin zymography. Supernatants were collected from overnight cultures with the same cell number. Higher protease activities correspond to clearer bands on the gel matrix. M, marker; Δ7, mutant lacking seven extracellular proteases (*ΔaprE, ΔnprE, ΔnprB, Δbpr, Δepr, Δmpr, Δvpr*).

To determine the origins of this small colony phenotype, we compared growth of WT and FY mutant strains on glucose-minimal medium (MM) agar plates amended with the individual ingredients of LB medium. The FY mutant was most defective (∼67% decrease in colony area) in accessing the nutrients supplied as a tryptone (Fig. 1b), an enzymatic digest of casein. Thus, we hypothesized that FY is defective in secretion of proteases required to access oligopeptides^27^. Over half of the ∼50 mM of amino acids in LB medium are only detected after acid hydrolysis, consistent with the idea that they are present in oligopeptides^28^.

*B. subtilis* is widely appreciated for secreting proteins during transition and stationary phases^29^, including industrially relevant proteases^30^. By monitoring extracellular protease production on milk agar plates^31^, we observed halo formation up to a 10^-4^ dilution for WT, but only up to 10^-1^ dilution for FY (Fig. 1c). *B. subtilis* encodes seven major extracellular proteases (AprE, NprB, NprE, Epr, Bpr, Vpr and Mpr) with ∼95% of activity attributed to the serine protease subtilisin (AprE) and the major metalloprotease NprE^29^. We used zymography to determine which extracellular proteases might be deficient in the FY mutant. The reduction in protease levels in FY mutant was not restricted to the three metalloproteases with predicted molecular weights of 59 kDa (NprB), 56 kDa (NprE), and 34 kDa (Mpr) (Fig. 1d), and included a reduction in bands representing degradation products of the large Bpr protease (154 kDa)^32^ (Fig. S2). This overall reduction in protease activity led us to hypothesize that the FY mutant has a generalized defect in protein secretion.

### The FY mutant is defective in protein secretion

The FY mutant secretes ∼30% less protein than WT after overnight growth in LB (Fig. 2a). Since MeeF and MeeY have Mn export activity^20^, we hypothesized that they might play a specific role in secretion of Mn-requiring proteins. Consistently, the FY mutant had a >5-fold reduction in extracellular Mn (Fig. 2b). These results suggest that most Mn in the growth medium is imported to support growth. Indeed, our LB medium contains just enough Mn (126 nM) to support cell growth without limitation^25^ and more than 90% is used by cells during growth (FY mutants have ∼3.2 nM residual Mn in the spent medium, Fig. 2b). Thus, the higher level of Mn (∼22 nM) detected in the supernatant of WT (and single F and Y mutants) is likely associated with secreted metalloproteins. In contrast, these strains had little difference in the residual level of Fe or Zn detected in the spent medium (Fig. S3).

**Figure 2.**
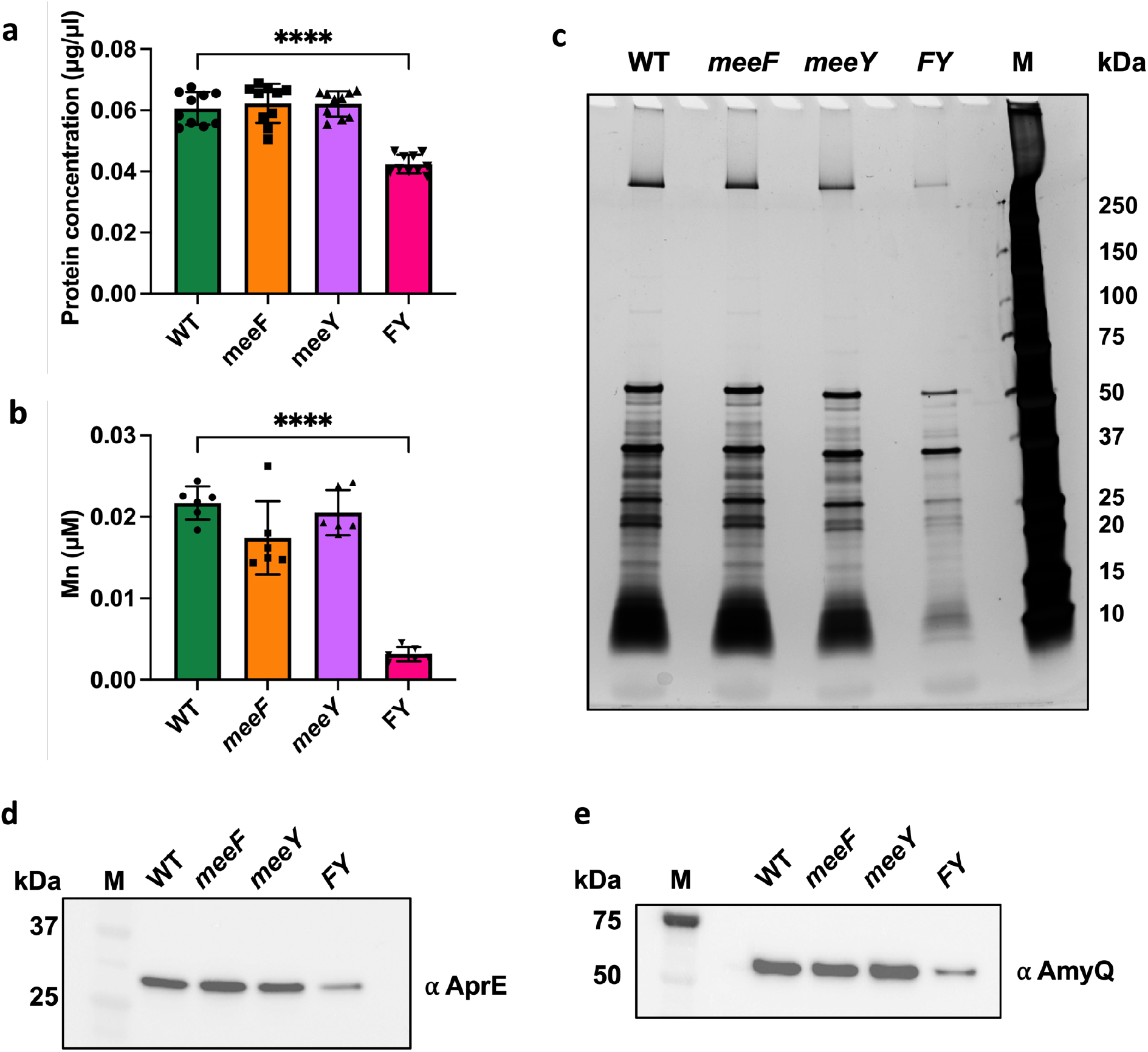
FY mutants have a generalized secretion defect. (**a**) FY mutants (but not the single F and Y mutants) have reduced levels of secreted proteins in the spent medium (supernatant fraction) after overnight culture. (**b**) FY mutants also have dramatically reduced levels of Mn in the spent medium after overnight growth as monitored by ICP-MS analysis. For (**a**) and (**b**), samples were from three independent experiments. Data is presented as mean ± SD. ****, P < 0.0001, P value was calculated using Welch’s t test, two-tailed. (**c**) Silver staining showing reduced extracellular protein in the supernatant from the FY strain compared to WT and the single mutant strains after overnight growth (representing the same final culture density; Fig S1b). The image is representative of three experiments. (**d**) The FY mutant is defective for secretion of AprE-FLAG. (**e**) The FY mutant is defective for secretion of heterologous AmyQ-His protein. For (**d**) and (**e**), strains were grown overnight to same cell density and centrifuged to obtain supernatant and pellet fractions. The levels of proteins were probed by immunoblotting with anti-FLAG or anti-His antibodies. Representative images of three independent experiments are shown. The full gels and Ponceau-stained images (to demonstrate equal loading) are shown in extended data Fig. S4. M, all stain precision blue marker (BioRad).

Analysis of extracellular proteins revealed a general reduction in the supernatant of the FY strain (Fig. 2c). To further evaluate the secretion capacity of the FY mutant, we probed the extracellular levels of two well-studied secretion substrates: the major extracellular protease subtilisin E (AprE) and α-amylase from *B. amyloliquefaciens* (AmyQ)^29, 30, 33^. We observed a >2-fold decrease for both AprE-FLAG^34^ and α-amylase AmyQ-His^35^ in the FY mutant, but not in the F and Y single mutants (Fig. 2d, 2e, S4). Therefore, MeeF and MeeY have overlapping functions required for efficient protein secretion.

The high-level overproduction of secreted proteins such as AmyQ can lead to activation of the secretion stress response controlled by the CssRS two-component system^29^. CssS senses accumulation of misfolded exoproteins to activate expression of the quality control proteases HtrA and HtrB^36, 37^. Consistent with their reduced secretion capacity, FY mutants do not experience secretion stress, as monitored using the CssR-dependent P*_htrA-lux_* reporter. However, the P*_htrA-lux_* reporter was still induced upon overexpression of AmyQ (Fig. S5), despite reduced AmyQ export (Fig. 2e). Indeed, the FY mutant had a slightly increased P*_htrA-lux_* induction relative to WT (Fig. S5), similar to other secretion-deficient mutants^38^.

### MeeF and MeeY interact with the general secretory pathway

Most membrane and secreted proteins are translocated by the general secretion pathway (Sec pathway) in *B. subtilis*^39^. To investigate how MeeF and MeeY might influence secretion efficiency, we used co-immunoprecipitation to identify proteins that interact with C-terminal FLAG-tagged MeeF or MeeY. Expression of both proteins was increased in LB medium supplemented with 50 μM Mn, and MeeY-FLAG had elevated expression in a mutant lacking MeeF (Fig. S6a). After co-immunoprecipitation, shotgun proteomics was used to identify co-purifying proteins (Table 1, Fig. S6b, S6c). We focused on those putative interactors that are membrane proteins, since both MeeF and MeeY are integral membrane proteins largely comprising 7 transmembrane segments. Strikingly, many of the co-purifying proteins (Table 1) were either components of the holotranslocon (SecDF, SecY, YrbF), quality control proteases (FtsH, PrsA, HtpX), or subunits of the F_1_F_o_ ATPase, which forms a complex with the SecYEG translocon^39–42^. Thus, MeeF and MeeY appear to function as part of the secretosome, likely by mediating co-translocational protein metalation.

**Table 1.** Membrane Proteins that Co-immunoprecipitate with MeeF and MeeY.

| MeeF-FLAG<br>(heating)<sup>a</sup> | MeeY-FLAG<br>(heating)<sup>a</sup> | MeeY-FLAG<br>(pH)<sup>b</sup> | Functional annotation |
| --- | --- | --- | --- |
| **Holotranslocon Proteins** |  |  |  |
| SecY |  |  | Subunit of the SecYEG preprotein translocase |
| SecDF | SecDF | SecDF | PMF-dependent holotranslocon subunit |
| YrbF |  |  | Binds to SecDF as part of the holotranslocon (*E. coli* YajC ortholog; 34% identity) |
| **Secretosome and Secretion-related Functions** |  |  |  |
| PrsA | PrsA | PrsA | post-translocation molecular chaperone |
| HtpX |  |  | Quality control membrane protease |
| FtsH | FtsH |  | Quality control membrane protease |
| AtpA | AtpA | AtpA | ATP synthase (subunit alpha); intact ATP synthase interacts with SecYEG translocon<sup>16</sup> |
| AtpG | AtpG |  | ATP synthase (gamma subunit) |
|  | AtpF |  | ATP synthase, part of the Fo complex |
|  |  | AtpD | ATP synthase, part of the F1 complex (subunit beta) |
| **Other Membrane Proteins** |  |  |  |
| FloT |  |  | Flotillin, membrane-associated scaffold protein |
| FloA | FloA |  | Flotillin, membrane-associated scaffold protein |
| MalA |  |  | 6-phospho-alpha-glucosidase |
| GlcD |  |  | possible glycolate oxidase subunit |
| NupN |  |  | lipoprotein, part of guanosine transporter |
| SwrC | SwrC |  | resistance-nodulation-cell division (RND) |
| SsdC |  |  | spore shape determinant C (mother cell) |
| OppF | OppF |  | Oligopeptide ABC transporter |
| OppB |  |  | Oligopeptide ABC transporter |
| FrlO |  |  | Aminosugar ABC transporter |
| SdhA | SdhA |  | succinate dehydrogenase (flavoprotein subunit) |
| FhuD |  |  | hydroxamate siderophore ABC transporter |
|  | QoxA |  | cytochrome aa3 quinol oxidase (subunit II) |
| QoxB |  |  | cytochrome aa3 quinol oxidase (subunit I) |
| MsmX |  |  | multiple sugar ABC transporter (ATP-binding protein) |
| FtsA |  |  | cell division protein, member of the <u>divisome</u> |
| YknW | YknW |  | modulator of ABC transporter assembly, SdpC secretion |
|  |  | MgtE | primary magnesium transporter |

<sup>a</sup>Immunoprecipitated fractions were eluted by heating, samples were treated with 1% TritonX-100 and their western analysis is shown in Fig S6b.
<sup>b</sup>Immunoprecipitated fractions were eluted by glycine (pH 3), the MeeY-FLAG sample is shown in Fig S6c.

### FY mutants require the FtsH protease for viability

FtsH is an ATP-dependent metalloprotease that functions to degrade membrane proteins in response to a variety of stresses^21^. Studies in *E. coli* indicate that FtsH selectively targets membrane-associated proteins that are misfolded, including partially translocated proteins stalled during passage through the SecYEG translocon (translocon jamming)^43^. Indeed, FtsH can degrade the major translocase subunit SecY^44^ and this activity can even lead to a lethal defect in protein secretion if not properly regulated^45^. Phenotypically, *ftsH* mutants are similar to FY mutants in that they display a small colony size and are defective for protein secretion (Fig. S7)^46^.

We hypothesized that nascent metalloproteins might jam the SecYEG translocon in the FY mutant leading to the observed global impairment in protein secretion. Under this condition, the FtsH protein is predicted to play an important role in removing partially translocated proteins and in clearing jammed translocons from the membrane^15^. To test this idea, we attempted to construct an FY *ftsH* triple mutant strain. While it was possible to generate all three possible double mutants, the triple mutant was inviable and efforts to construct this strain by genetic transformation invariably led to congression (acquisition of a functional copy of one of the missing genes). Conversely, induction of FtsH helped rescue the poor growth phenotype of FY mutant cells (Fig. S7a). This is consistent with models that posit a role in FtsH in removal of partially translocated proteins from stalled translocons^15^. These genetic interactions support the hypothesis that MeeF and MeeY act in support of the translocation of nascent metalloproteins, and their absence leads to translocon jamming.

### MeeF and MeeY support activity of Mn-dependent lipoteichoic acid synthases

LtaS is the major lipoteichoic acid (LTA) synthase in *B. subtilis*^47, 48^. LtaS is an integral membrane protein with an extracytosolic globular domain that requires Mn for activity^48^. One phenotype of an *ltaS* mutant is small colony size ^49^, similar to FY (Fig. S7a). We therefore hypothesized that MeeF and MeeY may play a role in the loading of Mn into LtaS.

To determine if MeeF and MeeY have a role in the activation of LtaS, we used immunoblotting to monitor levels of LTA^50^. LTA levels were similar in the WT strain and the *meeF* single mutant, reduced in the *meeY* mutant, and greatly reduced in the FY double mutant (Fig 3a). The specificity of the assay is apparent from analysis of the *ltaS* single mutant, which lacks the abundant ∼10-15 kDa LTA polymers. In this strain, longer LTA chains were produced that are the product of the stress-induced alternate LtaS enzyme, LtaSa(YfnI)^48^, as confirmed by their absence in the *ltaS ltaSa* double mutant (Fig. 3a). These results indicate that either MeeF or MeeY can support LtaS function.

**Figure 3.**
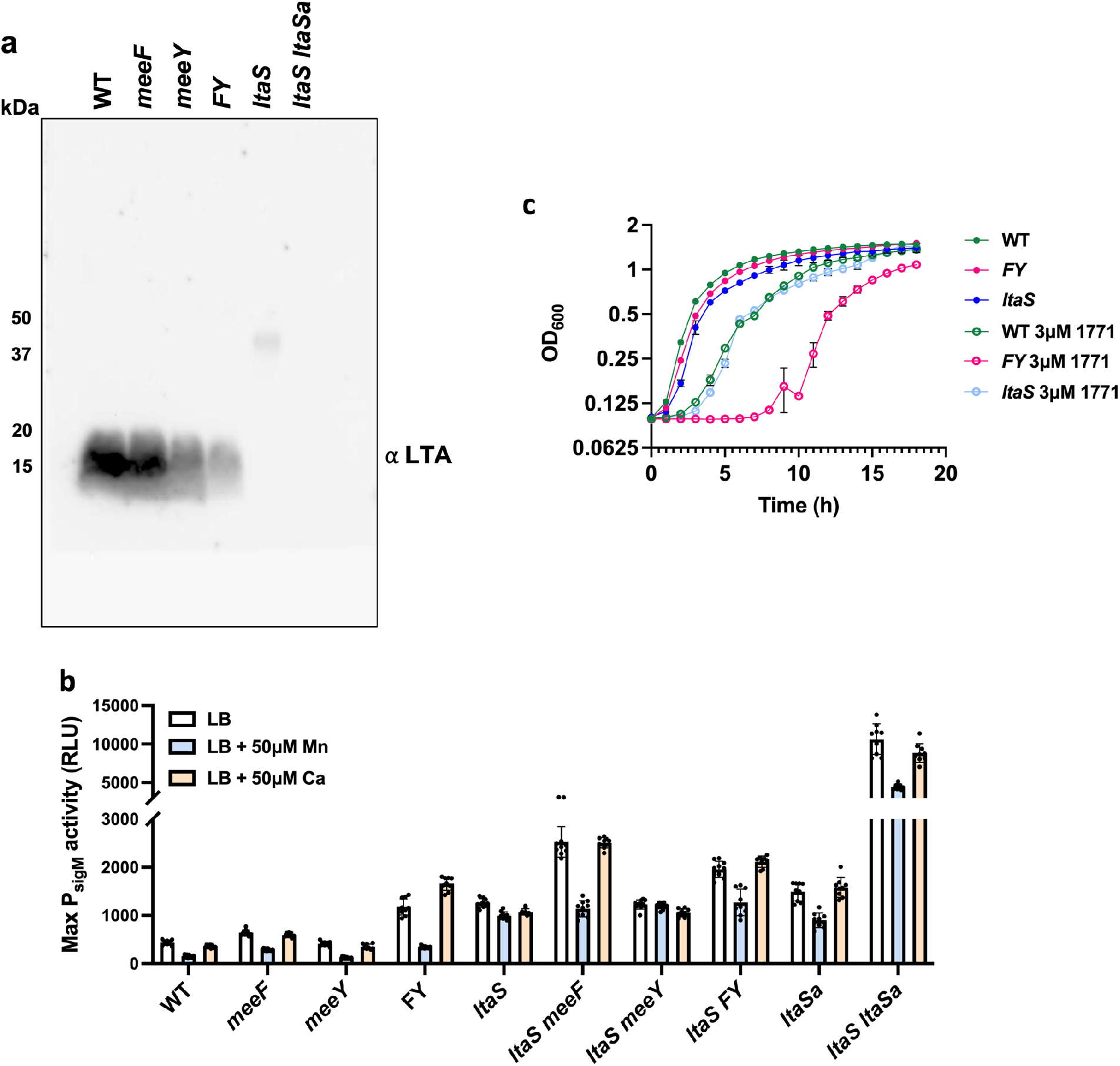
FY mutants are defective in LTA synthesis. (**a**) Immunoblot detection of LTA with anti-LTA monoclonal antibodies. Note that in *ltaS* mutants the signal in the ∼15-20 kDa range is absent, and instead longer polymers are detected that depend on the LtaSa enzyme^48^. The image is representative of two independent experiments with gels loaded with extracts from equal cell numbers. (**b**) Defects in LTA synthesis activate the α^M^-dependent envelope stress response as monitored using a luciferase transcriptional reporter fusion (P*sigM-luxABCD*). Cells were grown in LB broth with or without Mn (50 µM) or Ca (50 µM). Data is from three independent biological experiments and shown with mean ± SD. (**c**) Defective metalation of LTA synthase enzymes is associated with increased sensitivity to compound 1771, a LtaS inhibitor^54, 87^ . Aerobic growth of different strains (WT, *FY, ltaS*) in LB broth with or without 3 µM 1771 is shown. Data are representative of three independent cultures and presented as mean ± SD. Additional results, showing the effects of metal supplementation are in Fig. S9.

Cells mutant for *ltaS* experience cell envelope stress, due in part to dysregulation of autolysins ^50^, and activate the α^M^ cell wall stress response^51^ to express an alternate LTA synthase, LtaSa ^52^. Since FY mutant cells are defective in LTA synthesis, we predicted that they would also activate the expression of the α^M^ stress response. Consistent with our hypothesis, the FY double mutant (but not F or Y) had elevated expression of a σ^M^-dependent promoter (P*_sigM-lux_*) ^53^ (Fig. 3b). The level of activation of σ^M^ in the FY mutant was comparable to that seen in a strain lacking *ltaS*. Further, σ^M^ activation was even higher in *ltaS* strains additionally lacking *meeF (ltaS meeF)*, but not in *ltaS meeY* double mutants (Fig. 3b). This additivity suggests that cell stress was increased by mutation of MeeF in strains lacking the major LtaS enzyme. Thus, LtaSa may also require Mn. Indeed, prior results demonstrate that *ltaS ltaSa* double mutants have an elevated stress response ^51^, as also seen here (Fig. 3b). In addition to LtaS and the stress-inducible synthase LtaSa, *B. subtilis* expresses YqgS (a minor LTA synthase) and YvgJ (an LTA primase)^48^. We monitored the expression of all four LTA synthesis genes using quantitative RT-PCR. As expected, the level of *ltaSa* mRNA was elevated in the FY mutant, but not in the single mutant strains (Fig. S8).

Since the catalytic domain of LtaS is external to the cell, we next tested whether addition of Mn to the growth medium could activate enzyme that had been properly inserted in the membrane but had failed to acquire its catalytic Mn ion. Indeed, Mn reversed the σ^M^ cell envelope stress response in the FY mutant but not, as expected, in the *ltaS* mutant (Fig. 3b). Addition of Ca, which is not an effective co-factor for LtaS enzymes^48^, did not reverse induction of the σ^M^ stress response. Thus, the FY mutant has properly expressed LtaS, inserted it in the membrane, it failed to acquire Mn. Addition of Mn also partially reversed the σ^M^ stress response seen in the *ltaS meeF* mutant (and *ltaS FY* mutant), consistent with the hypothesis that this strain may be partially deficient in activation of LtaSa or other back-up synthases. The elevated stress response seen in *ltaS ltaSa* (Fig. 3b) was also partially reduced by Mn, possibly due to activation of the YqgS synthase.

### FY mutants are sensitized to chemical inhibition of LtaS activity

Because of its important role in physiology, LTA has been a target for the development new antibacterials^54^. An inhibitory compound LtaS-IN-1(1771) binds to the active site of LtaS^55, 56^. Binding of 1771 inhibits the interaction between LtaS and its substrate phosphatidylglycerol^54^. The FY mutant was much more sensitive to growth inhibition by 3 μM 1771 than WT (Fig. 3c). Further, *ltaS* mutant strains were as sensitive to growth inhibition by 3 μM 1771 as WT strain, presumably because 1771 is active against alternative LtaS enzymes (Fig. 3c). We therefore hypothesized that FY is more vulnerable to 1771 because it is deficient in Mn loading in the active sites of LtaS and LtaSa enzymes. To explore this hypothesis, we tested the effect of metal ion supplementation on sensitivity to 1771 in various mutant strains. In the presence of 1771, Mn improved growth of WT and FY mutant cells, but only slightly rescued the *ltaS* mutant (Fig. S9). In contrast, Zn worsened the 1771 growth inhibition in all tested strains, in some cases dramatically (Fig. S9). These results suggests that LTA synthases may be subject to mismetalation by Zn, and this inhibition is enhanced in the FY mutant where Mn acquisition has been compromised. Collectively, these results indicate that 1771 may bind to the active site of LtaS and its paralogs, and this may hinder or preclude activation under Mn-limiting conditions.

### The function of TerC proteins is conserved in Gram-positive bacteria

TerC proteins are conserved among different bacterial species^57^. To determine whether heterologous TerC proteins can complement the FY mutant we expressed two *Listeria monocytogenes* TerC proteins (Lmo0991, Lmo0992) from and one *B. anthracis* TerC (BanTerC). As expected, expression of either of the native TerC proteins (MeeF and MeeY) restored colony size on LB, and similar results were seen with heterologous TerC proteins (Fig. 4a). Consistently, the protease activity of these complemented strains was also restored (Fig. 4b). Therefore, the function of TerC proteins is conserved across Gram-positive microbes.

**Figure 4.**
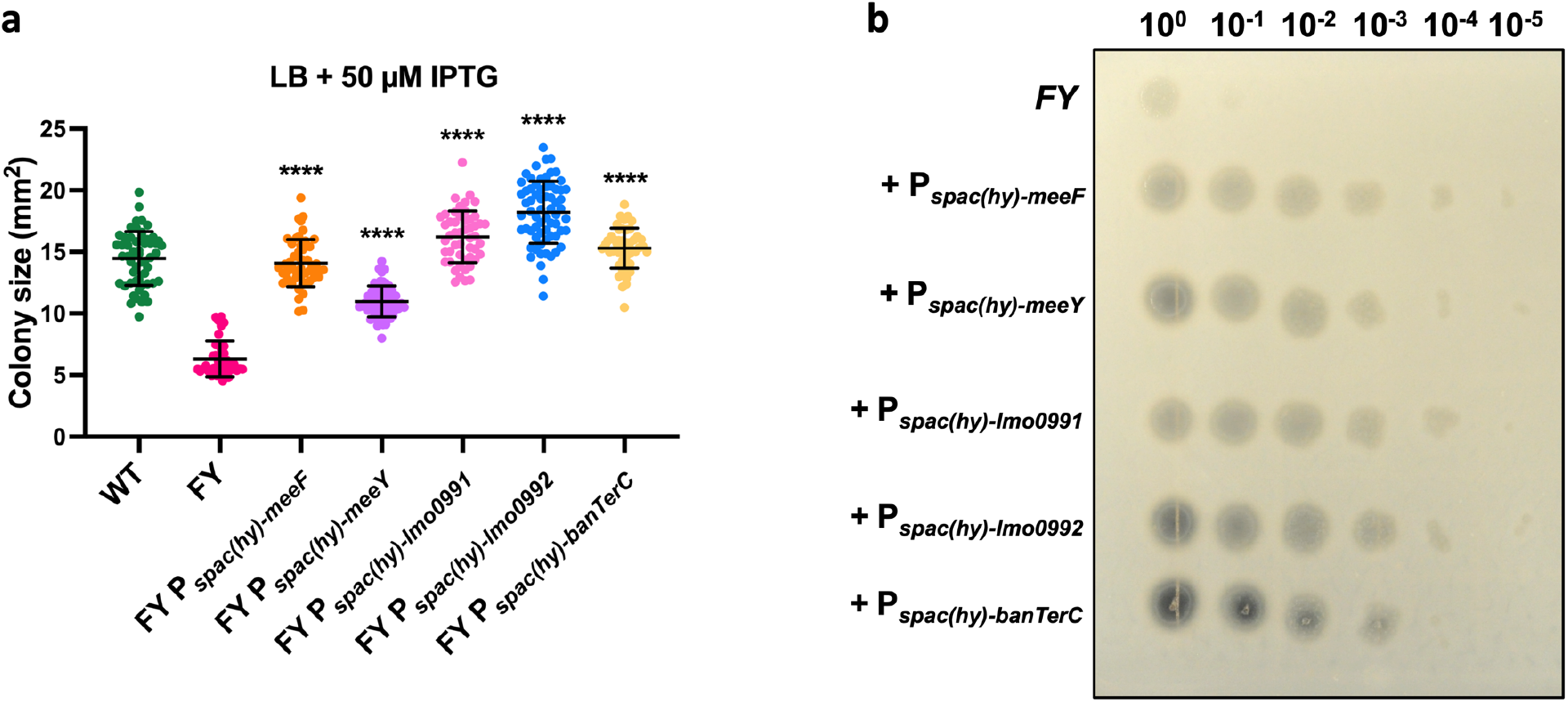
Complementation of the FY mutant with orthologous TerC proteins. (**a**) Colony size (mean ± SD) of FY mutant with induction of TerC proteins on LB medium with 50 µM IPTG. TerC proteins were induced by 50 µM IPTG using Pspac(hy) promoter. Colony size was measured by imageJ. At least 40 isolated colonies measured for each strain. ****, P<0.0001, P value of each strain compared to FY samples was calculated using Welch’s t test, two-tailed. (**b**) Protease activities of FY mutant and FY complementary strains on 5% milk agar plates are shown. Cells were grown in LB broth with 50 µM IPTG inducer to OD600 0.4. Serial diluted cells (10^0^ – 10^-5^) from these cultures were inoculated on the plates 37°C for 24 hours.

## Discussion

The metabolic processes that support life would grind to a halt without the catalytic enhancements enabled by metal ions^2^. The most widely deployed metal ions in cytosolic enzymes are Zn, Fe, and Mn, with Cu enzymes largely restricted to the membrane^1^. Most enzymes are active only when associated with the correct metal and have therefore evolved metal-binding sites that confer selectivity for the relevant metal ion^4, 5^. However, in some cases alternative metals may also sustain activity. Conversely, binding of the wrong metal (mismetalation) may lead to enzyme inactivation^58^. To ensure proper metalation, Cu enzymes may require metallochaperones^59, 60^. Similarly, metallochaperones have been identified for selected Zn enzymes in bacterial^61, 62^, fungal^63^, and mammalian cells^64^.

How proteins that are exported from the cell acquire the proper metal ion is less well understood. For high affinity metals from the Irving-Williams series, acquisition from the environment may suffice to ensure metalation. This is the likely route for Cu acquisition by the abundant periplasmic CucA(Cu-cupin A) protein from *Synechocystis* PCC 6803^9^ and for Zn-beta-lactamases^10^. For lower affinity metals, alternative strategies may be needed. For example, the MncA(Mn-cupin A) from *Synechocystis* PCC 6803 is metalated inside the cell and the folded protein exported through the TAT system^9^.

TerC proteins (Pfam03741) are a subgroup of the LysE superfamily of transporters with seven TM segments and a conserved metal-binding site^19, 65^. Although originally implicated in resistance to toxic tellurite salts^57^, there is no evidence that TerC proteins transport tellurite ^66, 67^. We identified up-regulation of *meeF* in a screen for suppressors of the high Mn sensitivity of strains lacking the MneP and MneS Mn efflux proteins^25^, and found that both MeeF and its MeeY have Mn efflux activity^20^. The three *B. subtilis* TerC proteins (MeeF, MeeY, and YjbE) are differentially regulated: *meeY* gene is regulated by a Mn-responsive riboswitch^68–70^, *meeF* is part of the constitutively expressed *yceCDEmeeF* operon and can be further induced by stress-responsive sigma factors^52, 71, 72^, and *yjbE* is expressed during sporulation^73^.

Here, we have identified MeeF and MeeY as accessory subunits of the holotranslocon that mediate the metalation of exoenzymes. Strains lacking both MeeF and MeeY (FY mutants) are defective in protein secretion, have a greatly reduced level of Mn in the cell supernatant, and fail to efficiently metalate LTA synthases. Co-immunoprecipitation reveals an association with proteins in the holotranslocon and secretosome complex. Further, FY mutants have strong epistatic interactions with *ftsH*, encoding a quality control protease that helps rescue jammed translocons. The most parsimonious explanation of these data is that TerC proteins function as metallochaperones to load Mn into nascent metalloproteins. However, they may instead, or in addition, simply export Mn to generate a sufficiently high local concentration to ensure metalation of proteins during secretion (Fig. 5). In their absence, exoenzymes may be deficient in metalation, and nascent metalloproteins may contribute to translocon-jamming.

**Figure 5.**
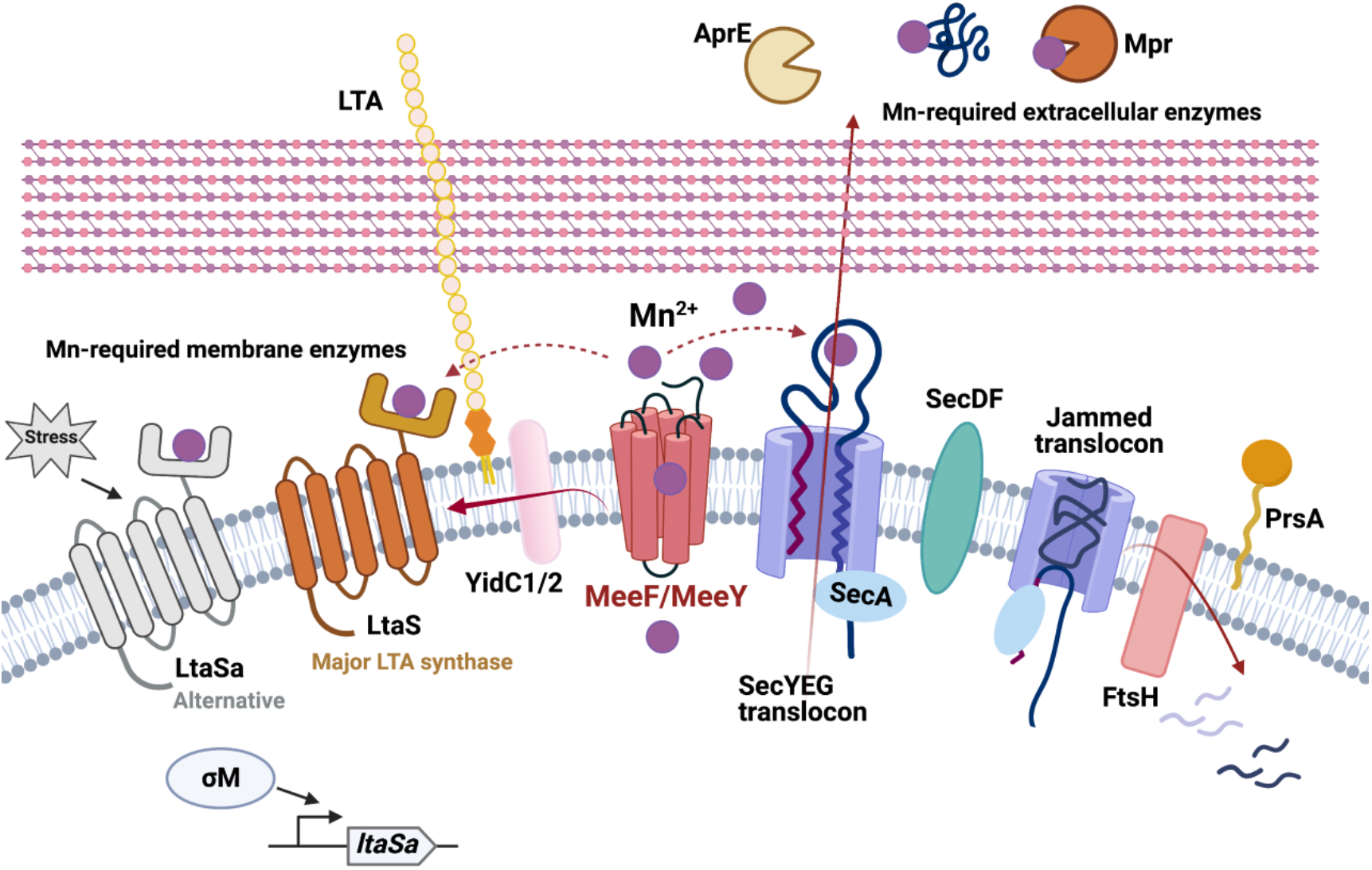
The functions of TerC proteins MeeF and MeeY in exoenzyme metalation. MeeF(F) and MeeY(Y) are integral membrane proteins that function in Mn export^20^. F and Y are here shown exporting Mn to support metalation of exoenzymes. F and Y interact physically (co-immunoprecipitation) and genetically (epistasis with *ftsH*) with proteins of the secretosome complex. These results suggest that F and Y function co-translocationally to insert Mn into nascent metalloproteins. As a result, FY double mutants are deficient in Sec-dependent secretion of exoenzymes (e.g. proteases, AprE, AmyQ), which leads to growth defects on LB medium. FY mutants are also deficient in activation of LTA synthases, which bind Mn to an extracellular domain to catalyze lipoteichoic acid synthesis. The essentiality of FtsH in the FY mutant is consistent with jamming of the SecYEG translocon. F and Y may function as metallochaperones that directly transfer Mn to client proteins and/or they may help generate a sufficiently high local Mn concentration to allow metalation.

Functional studies have linked diverse TerC proteins to the transport of Mn and Ca. In plants, *Arabidopsis thaliana* (AtTerC) is important for the insertion of thylakoid membrane proteins and interacts with the membrane protein insertase ALB3, a YidC ortholog^22, 74^. In yeast, Gdt1p mediates Mn influx into the Golgi to activate metalloenzymes functioning in protein glycosylation^75–77^. The human ortholog TMEM165 has a similar role as Gdt1p and missense mutations are associated with congenital disorders of glycosylation (CDG)^75, 78, 79^. These results suggest that TerC proteins are important in both bacteria and eukaryotes for the proper functioning of exported proteins, possibly by mediating the co-translocational insertion of metals into nascent proteins during transit across membranes.

## Online Methods

### Bacterial strains and growth conditions

All strains used in this study are listed in Table S1. Mutant strains were obtained from the *Bacillus* Genetic Stock Center (BGSC) as erythromycin marked gene disruptants from the BKE collection^80^. Mutations were transformed into the desired strain and markerless in-frame mutants were generated by transformation with plasmid pDR244 to remove the erythromycin cassette^80^. Gene deletions were confirmed by PCR screening using flanking or internal primers (Table S1). The AmyQ-His overexpression plasmid pKTH10^35^ was selected with15 μg/ml kanamycin.

For construction of FLAG-tagged gene fusions we PCR amplified the C-terminal ∼500-700 bp of the *meeF, meeY* and *aprE* genes with primers (Table S1). The PCR products were restriction digested and ligated into pre-digested pMUTIN-FLAG^81^ plasmid using T4 DNA ligase (NEB). The constructs were transformed into *E. coli* DH5α and TG1 strains selected with ampicillin (100 μg/ml). The recombinant plasmids were transformed into *B. subtilis* and integrated into the chromosome with erythromycin (1 μg/ml) selection (Table S1). For IPTG-based Pspac(hy) overexpression construction, genes were amplified by PCR using high-fidelity Phusion polymerase and PCR products were digested by restriction enzymes (Xbal and BglII) and ligated to pPL82^82^.The ligation products were transformed into *E. coli* DH5α. Then constructs were integrated into *amyE* and selected with chloramphenicol (10 μg/ml).

### Growth conditions

Bacteria were grown in liquid or solid lysogeny broth (LB) (Affymetrix) at 37°C unless otherwise stated. LB medium contains 10 g tryptone, 5 g yeast extract, and 5 g NaCl per liter. Antibiotics used for selecting *B. subtilis* strains include: spectinomycin 100 μg/ml, macrolide-lincosamide-streptogramin B (MLS = 1 μg/ml erythromycin + 25 μg/ml lincomycin), kanamycin 15 μg/ml, and chloramphenicol 10 μg/ml. Glucose minimal medium (MM) was prepared as a 2X MM stock made using a 10X Bacillus salt solution [(NH_4_)_2_SO_4_ 20 g/L, Na_3_C_6_H_5_O_7_·2H_2_O 10 g/L, L-glutamic acid potassium salt monohydrate 10 g/L], and adding 80 mM MOPS buffer (pH 7.4 using KOH), 4 mM KPO_4_ Buffer (pH 7.0), 20 µg/L tryptophan, 2% glucose, 160 µM MnCl_2_, 1.6 mM MgSO_4_, 8.8 mg/L ferric ammonium citrate. For pouring plates, equal volumes of filter-sterilized 2X MM stock and 3% autoclaved agar were mixed. 1% Tryptone, 1% NaCl or 0.5% Yeast Extract (Y.E.) were added into MM where indicated.

### Colony size measurements

Bacterial cells were grown in liquid LB medium or liquid minimal medium at 37°C with vigorous shaking to mid-exponential phase (OD_600_∼0.4-0.5), serially diluted, and plated onto 15 ml fresh LB or MM agar plates with different amendments as noted. Plates were incubated at 37°C for 24 hours prior to imaging. Colony size was measured using Fiji ImageJ per software’s instruction^83^.

### Protease activity on skim milk agar

Casein degradation in skimmed milk agar plates (5% skim milk and 1% agar) was used to assess protease activity by formation of a clear zone. Bacterial cells were grown in liquid LB medium at 37°C with vigorous shaking to mid-exponential phase (OD_600_∼0.4). Cultures of identical OD were serially diluted from 10^0^ to 10^-5^. 2 µl of cells were inoculated on the plates. Plates were incubated at 37°C for 24 h and then imaged.

### Proteolytic profile by Zymography

Zymography was performed as described previously^84^ and proteases assigned as described^32^. The thickness of Zymography gel was 1.5 mm, and the resolving gel contained 10% gelatin. Supernatants of 1 ml of overnight grown cultures were centrifuged at 15,000 g for 10 minutes, and then mixed with 2X sample buffer without reducing agent incubated at 37°C for 30 minutes. After electrophoresis, the gels were placed in the renaturing buffer (2.5% Triton X-100) and incubated at room temperature for 30 minutes without shaking. Then the gels were placed into activation buffer (50 mM Tris-HCl, pH 7.5, 1% Triton X-100 and 25 mM CaCl_2_) and incubated at 37°C for 18 hours. The gels were stained by Coomassie blue for 2 hours and destained overnight (10% acetic acid, 40% methanol). When the desired pale bands were observed in the gels, images were taken. Note that this assay is selective for those proteases that are easily renatured following SDS-PAGE and have activity with gelatin^84^.

### Metal ion quantification by ICP-MS

Metal content of supernatant fractions was measured as described previously^85^. Cells were grown overnight in LB medium, and the supernatants obtained by centrifuging 2 ml of the cultures at 15,000 g for 10 minutes. The protein concentrations of the supernatants were measured using the Bradford assay (Bio-Rad, USA) using BSA as a standard. 900 µl of supernatants were mixed with 600 µl of buffer (5% HNO_3_, 0.1% Triton X-100) and incubated at 95°C for 30 minutes. After centrifuging the samples at 15,000 g for 10 minutes, 1 ml of clear supernatants were transferred to new tubes, and the total metal ions in the supernatants were analyzed using a Perkin-Elmer Elan DRC II ICP-MS.

### Silver stain for protein detection in polyacrylamide gels

Cells were grown in LB liquid medium overnight at 37°C. Supernatants were obtained by spinning down 2 ml of the cultures at 15,000 g for 10 minutes and filtered using a low protein-binding polyethersulfone (PES) membrane sterile filter (Foxx Life Sciences). 500 µl of the supernatants were then mixed with 2X denaturing Laemmli sample buffer (Bio-Rad, USA) and incubated at 95°C for 5 minutes. The samples were quickly centrifuged and 10 µl of each supernatant was loaded onto a 4-20% stain-free polyacrylamide gel (Bio-Rad, USA). After electrophoresis, the gel was stained as described in the Pierce silver stain kit (Thermo Scientific^TM^ 24600). Briefly, the gel was fixed by fixing solution (30% ethanol, 10% acetic acid). After ethanol wash and water wash, the gel was incubated in sensitizer working solution and then stain working solution. After adding developer working solution, the bands appear on the gel after 2-3 minutes. Images were taken after adding stop solution (5% acetic acid) using a GelDoc Gel imaging system (Bio-Rad, USA).

### Protein detection by immunoblot

Samples were collected from overnight cultures without or with 50 µM Mn. About 5 ml of the cultures of identical OD were centrifuged; pellets and supernatants were separately collected. Cells were resuspended in 100 µl lysis buffer (20 mM Tris-HCl pH8.0, 1 mM EDTA, 1 mg/ml lysozyme) at 37°C for 30 minutes. The crude cell lysates and supernatants were mixed with 2X denaturing Laemmli Sampler buffer (Bio-Rad, USA) and boiled at 95°C for 10 minutes. The samples were cooled and fast centrifuged. 12 µl of samples were electrophoresed onto a 4-20% strain-free polyacrylamide gel (Bio-Rad). Proteins were then transfer onto a PVDF membrane using the Trans-Blot Turbo Transfer System (Bio-Rad, USA). The PVDF membrane was stained by Ponceau dye (5% glacial acetic acid, 0.1% ponceau S tetrasodium salt) for 15 minutes. The image was taken after removing Ponceau dye and rinsing the membrane with TTBS (1X TBS with 0.1% Triton X-100). The Ponceau stain was removed by repeated washing with TTBS, and the membrane was blocked with 5% protein blotting blocker dissolved in TTBS for 1 hour. The primary antibodies, anti-FLAG antibody produced in rabbit (Sigma); or 6X His Tag antibody in mouse (Invitrogen) were added in 0.5% protein blotting blocker dissolved in TTBS (1:5000), and the membrane was incubated with the primary antibodies overnight at room temperature. After washing the membrane three times with TTBS for 10 minutes, the membrane was incubated in the secondary antibodies, Mouse anti-rabbit HRP (Cell Signaling Technology); or Rabbit anti-Mouse secondary antibody, HRP (Invitrogen) for 90 minutes (1:10000). The membrane was washed five times with TTBS and then visualized using the Clarity Western ECL substrate (Bio-Rad, USA). Band intensity was calculated using ImageJ software.

### LTA detection by Western blot

Samples were collected as described^50^. Stains were grown in 5 ml of nutrient broth (NB) with 5 mM MgSO_4_ and shaking at 30°C overnight. Cultures were diluted to 0.01 OD_600_ in NB and then shaking at 37°C for about 5 hours. Cultures with identical cell numbers (OD_600_∼0.6) were collected. Sample collection and LTA detection by Western blot was same as described^50^. We used 2X Laemmli Sampler buffer instead of lithium dodecyl sulfate to resuspend samples.

### Co-immunoprecipitation (Co-IP) of MeeF-FLAG and MeeY-FLAG

Cells were collected from 10 ml of overnight LB cultures containing 50 µM Mn and were lysed by 1 ml of lysis buffer (Tris-HCl, 50 mM EDTA, 1 mg/ml lysozyme) incubated at 37°C for 30 minutes. The samples were sonicated for 5 minutes and incubated with 10 U/ml DNase at 37°C for 30 minutes. 1% Triton X-100 was added, and then samples were incubated on ice for 2 hours before adding anti-FLAG M2 magnetic beads (Sigma). Co-IP samples were prepared as described in the protocol of the anti-FLAG M2 magnetic beads (Millipore Sigma). Samples were end-over-end rotated with anti-FLAG M2 magnetic beads at 30°C for 3 hours and then placed on a magnetic stand.

Supernatants were discarded and the samples were washed three times with PBS buffer at 30°C for 10 min. Samples were eluted be either heating at 95°C for 10 minutes or treatment with glycine (pH 3) for 30 minutes at room temperature. Proteins in the samples were checked by immunoblot and were identified by LC-MS/MS (Cornell Proteomics and Metabolomics Facility). Only proteins with at least 2 peptides were considered as positive identifications. Table 1 summarizes all membrane-localized proteins from three experiments.

### Promoter-luciferase measurements

The activity measurements of promoter P*_htrA_* and P*_sigM_* were as described^86^. Bacterial cells were grown aerobically in liquid LB medium at 37°C to OD_600_∼0.4. 1 μl of the cultures were inoculated into 99 μl of fresh liquid LB without or with different metals dispensed in a 96-well plate. The plate was incubated at 37°C with vigorous shaking using a Synergy H1 (BioTek Instruments, Inc. VT) plate reader. OD_600_ and luminescence were measured every 6 minutes. The maximum promoter activity was measured by dividing the relative light units (RLU) with OD_600_.

### Real Time RT PCR

Total RNA was extracted using QIAGEN kit from 1.5 ml of mid-log (OD 600nm =0.4) WT, *meeF*, *meeY*, and FY bacterial cultures, which were grown in LB broth either in the presence or absence of 0.1 mM Mn. Total RNA was treated with DNase (Ambion) enzyme to further purify and remove traces of DNA. For each reaction 2 µg was used for cDNA synthesis using High-Capacity reverse transcriptase (Applied Biosystems) amplified with random hexamer primers. Further, for amplicon measurements 10 ng of cDNA was used as a template along with 500 nM of *ltaS*, *ltaSa(yfnI)*, *yqgS*, *yvgJ* and *gyrA* (control) gene specific qPCR F/R primers in a 1X SYBR green master mix (Bio-Rad). Threshold and baselines parameters were kept consistent for experiments performed on a different day during data analysis. All CT mean values for the gene expressions were normalized to *gyrA* (n=2).

### Growth curve with LtaS inhibitor 1771

Bacterial cells were grown in liquid LB medium at 37°C to OD_600_∼0.4. 2 μl of the cultures were inoculated into 198 μl of fresh liquid LB with different concentrations of 1771 (MedChemExpress) and dispensed in a 96-well plate. The plate was incubated at 37°C with vigorous shaking using a Synergy H1 (BioTek Instruments, Inc. VT) plate reader. OD_600_ was measured every 30 minutes.

## ACKNOWLEDGEMENTS

This work was supported by National Institutes of Health R35GM122461 awarded to JDH. We acknowledge Sri Paruthiyil for his initial observation of the small colony phenotype on LB medium, Brian Wendel for help with ICP-MS analyses, the Cornell Biotechnology Resource center for proteomics analysis, and Prof. Jan Maarten Van Dijl for the gift of pKTH10 plasmid.

## Author Contributions

B.H., A.J.S., and J.D.H designed the experiments. B.H. and A.J.S. performed the experiments. B.H, A.J.S., and J.D.H wrote the paper.

