## Supplemental Files for "TerC Proteins Function During Protein Secretion to Metalate Exoenzymes"

**Table S1**. *Bacillus subtilis* stains, primers, and plasmids used in this study.

**Figure S1**. Comparative growth of WT and FY mutants.

**Figure S2**. Extracellular protease activities in the supernatants of protease mutants.

**Figure S3**. Metal ion levels (µM) measured in cell supernatant fractions.

**Figure S4**. Effect of *meeF*, *meeY* and FY mutations on AprE and AmyQ secretion.

**Figure S5**. Maximum $P_{htrA}$-lux promoter activity in stains with or without AmyQ overexpression

**Figure S6**. MeeF-FLAG and MeeY-FLAG in different mutants.

**Figure S7**. Comparative phenotypes of mutations affecting protein secretion.

**Figure S8**. Fold change of transcription level of LTA synthase genes in different strains.

**Figure S9**. Effects of metal ions of sensitivity to the LtaS inhibitor 1771.

**Table S1. Bacillus subtilis stains, primers, and plasmids used in this study.**

| Strain | Genotype | Construction | Reference |
|---|---|---|---|
| HB27501 | *trpC2 attSPβ (WT) CU1065* | Lab strain | Lab stock |
| HB27502 | *CU1065 ΔmeeF* | Lab strain | [1] |
| HB27503 | *trpC2 ΔmeeY* | Lab strain | [1] |
| HB27504 | *trpC2 ΔmeeF ΔmeeY* | Lab strain | [1] |
| HB27531 | *trpC2 aprE::erm* | BGSC-->HB27501 | This work |
| HB27534 | *trpC2 nprB::erm* | BGSC-->HB27501 | This work |
| HB27536 | *trpC2 nprE::erm* | BGSC-->HB27501 | This work |
| HB27541 | *trpC2 mpr::erm* | BGSC-->HB27501 | This work |
| HB27560 | *ΔnprE ΔaprE Δepr Δmpr ΔnprB Δvpr Δbpr* | | BGSC1A1133 |
| HB27584 | *trpC2 ftsH::erm* | BGSC-->HB27501 | This work |
| HB27637 | *trpC2 bpr::erm* | BGSC-->HB27501 | This work |
| HB27641 | *trpC2 epr::erm* | BGSC-->HB27501 | This work |
| HB27667 | *trpC2 ltaS:erm* | BGSC-->HB27501 | This work |
| HB27668 | *trpC2 ΔmeeF ltaS:erm* | BGSC-->HB27502 | This work |
| HB27669 | *trpC2 ΔmeeY ltaS:erm* | BGSC-->HB27503 | This work |
| HB27670 | *trpC2 ΔFY ltaS:erm* | BGSC-->HB27504 | This work |
| HB27680 | *trpC2 sacA::*$P_{htrA}$*-lux-cat* | gDNA HB23650-->HB27501 | [2] |
| HB27681 | *trpC2 ΔmeeF sacA::*$P_{htrA}$*-lux-cat* | gDNA HB23650-->HB27502 | This work |
| HB27682 | *trpC2 ΔmeeY sacA::*$P_{htrA}$*-lux-cat* | gDNA HB23650-->HB27503 | This work |
| HB27683 | *trpC2 ΔFY sacA::*$P_{htrA}$*-lux-cat* | gDNA HB23650-->HB27504 | This work |
| HB27684 | *trpC2 amyQ-His::kan* | pKTH10-->HB27501 | This work |
| HB27685 | *trpC2 ΔmeeF amyQ-His::kan* | pKTH10 -->HB27502 | This work |
| HB27686 | *trpC2 ΔmeeY amyQ-His::kan* | pKTH10 -->HB27503 | This work |
| HB27687 | *trpC2 ΔFY amyQ-His::kan* | pKTH10[c]-->HB27504 | This work |
| HB27721 | *trpC2 aprE-FLAG::MLS* | *aprE*::pMUTIN-FLAG-->HB27501 | This work |
| HB27722 | *trpC2 ΔmeeF aprE-FLAG::MLS* | *aprE*::pMUTIN-FLAG-->HB27502 | This work |
| HB27723 | *trpC2 ΔmeeY aprE-FLAG::MLS* | *aprE*::pMUTIN-FLAG-->HB27503 | This work |
| HB27724 | *trpC2 ΔFY aprE-FLAG::MLS* | *aprE*::pMUTIN-FLAG-->HB27504 | This work |
| HB27729 | *trpC2 sacA::*$P_{htrA}$*-lux-cat amyQ-His::kan* | pKTH10 -->HB27680 | This work |
| HB27730 | *trpC2 ΔmeeF sacA::*$P_{htrA}$*-lux-cat amyQ-His::kan* | pKTH10 -->HB27681 | This work |
| HB27731 | *trpC2 ΔmeeY sacA::*$P_{htrA}$*-lux-cat amyQ-His::kan* | pKTH10 -->HB27682 | This work |
| HB27732 | *trpC2 ΔFY sacA::*$P_{htrA}$*-lux-cat amyQ-His::kan* | pKTH10 -->HB27683 | This work |
| HB27733 | *trpC2 sacA::*$P_{sigM}$*-lux-cat* | gDNA *sacA::*$P_{sigM}$*-lux*-->HB27501 | [3] |

| | | | |
|---|---|---|---|
| HB27734 | *trpC2 ΔmeeF sacA::*$P_{sigM}$*-lux-cat* | gDNA *sacA::*$P_{sigM}$*-lux*-->HB27502 | This work |
| HB27735 | *trpC2 ΔmeeY sacA::*$P_{sigM}$*-lux-cat* | gDNA *sacA::*$P_{sigM}$*-lux*-->HB27503 | This work |
| HB27736 | *trpC2 ΔFY sacA::*$P_{sigM}$*-lux-cat* | gDNA *sacA::*$P_{sigM}$*-lux*-->HB27504 | This work |
| HB27739 | *trpC2 ltaS::erm sacA::*$P_{sigM}$*-lux-cat* | gDNA *sacA::*$P_{sigM}$*-lux*-->HB27667 | This work |
| HB27740 | *trpC2 ltaS::erm ΔmeeF sacA::*$P_{sigM}$*-lux-cat* | gDNA *sacA::*$P_{sigM}$*-lux*-->HB27668 | This work |
| HB27741 | *trpC2 ltaS::erm ΔmeeY sacA::*$P_{sigM}$*-lux-cat* | gDNA *sacA::*$P_{sigM}$*-lux*-->HB27669 | This work |
| HB27742 | *trpC2 ltaS::erm ΔFY sacA::*$P_{sigM}$*-lux-cat* | gDNA *sacA::*$P_{sigM}$*-lux*-->HB27670 | This work |
| HB27744 | *trpC2 ltaSa::spec* | gDNA *ltaSa::spec*-->HB27501 | [4] |
| HB27745 | *trpC2 ΔmeeF ltaSa::spec* | gDNA HB27744-->HB27502 | This work |
| HB27746 | *trpC2 ΔmeeY ltaSa::spec* | gDNA HB27744-->HB27503 | This work |
| HB27747 | *trpC2 ΔFY ltaSa::spec* | gDNA HB27744-->HB27504 | This work |
| HB27750 | *trpC2 ltaS::erm ltaSa::spec* | gDNA HB27744-->HB27667 | This work |
| HB27751 | *trpC2 ΔmeeF ltaS::erm ltaSa::spec* | gDNA HB27744-->HB27668 | This work |
| HB27752 | *trpC2 ΔmeeY ltaS::erm ltaSa::spec* | gDNA HB27744-->HB27669 | This work |
| HB27753 | *trpC2 ΔFY ltaS::erm ltaSa::spec* | gDNA HB27744-->HB27670 | This work |
| HB27770 | *trpC2 amyE::*$P_{spac(hy)}$*-meeF-cat* | pPL82-*meeF*-->HB27501 | This work |
| HB27772 | *trpC2 amyE::*$P_{spac(hy)}$*-lmo0991-cat* | pPL82-lmo0991-->HB27501 | This work |
| HB27773 | *trpC2 amyE::*$P_{spac(hy)}$*-lmo0992-cat* | pPL82-lmo0992-->HB27501 | This work |
| HB27774 | *trpC2 amyE::*$P_{spac(hy)}$*-BanTerC-cat* | pPL82-lmoBanTerC-->HB27501 | This work |
| HB27783 | *trpC2 ΔFY amyE::*$P_{spac(hy)}$*-meeF-cat* | pPL82-*meeF*-->HB27504 | This work |
| HB27784 | *trpC2 ΔFY amyE::*$P_{spac(hy)}$*-lmo0991-cat* | pPL82-lmo0991-->HB27504 | This work |
| HB27785 | *trpC2 ΔFY amyE::*$P_{spac(hy)}$*-lmo0992-cat* | pPL82-lmo0992-->HB27504 | This work |
| HB27787 | *trpC2 ΔFY amyE::*$P_{spac(hy)}$*-BanTerC-cat* | pPL82-lmoBanTerC-->HB27504 | This work |
| HB27788 | *trpC2 ltaSa::spec sacA::*$P_{sigM}$*-lux-cat* | gDNA *sacA::*$P_{sigM}$*-lux*-->HB27744 | This work |
| HB27789 | *trpC2 ΔmeeF ltaSa::spec sacA::*$P_{sigM}$*-lux-cat* | gDNA *sacA::*$P_{sigM}$*-lux*-->HB27745 | This work |
| HB27790 | *trpC2 ΔmeeY ltaSa::spec sacA::*$P_{sigM}$*-lux-cat* | gDNA *sacA::*$P_{sigM}$*-lux*-->HB27746 | This work |
| HB27791 | *trpC2 ΔFY ltaSa::spec sacA::*$P_{sigM}$*-lux-cat* | gDNA *sacA::*$P_{sigM}$*-lux*-->HB27747 | This work |
| HB27792 | *trpC2 ltaS::erm ltaSa::spec sacA::*$P_{sigM}$*-lux-cat* | gDNA *sacA::*$P_{sigM}$*-lux*-->HB27750 | This work |
| HB27832 | *amyE::*$P_{spac(hy)}$*-meeY-cat* | pPL82-*meeY*-->HB27501 | This work |
| HB27834 | *trpC2 ΔmeeY amyE::*$P_{spac(hy)}$*-meeY-cat* | pPL82-*meeY*-->HB27503 | This work |
| HB27836 | *trpC2 ΔFY amyE::*$P_{spac(hy)}$*-meeY-cat* | pPL82-*meeY*-->HB27504 | This work |
| HBAS924 | *trpC2 meeF-FLAG::MLS* | pMUTIN-FLAG | This work |
| HBAS943 | *trpC2 meeY-FLAG::MLS* | pMUTIN-FLAG | This work |
| HBAS927 | *trpC2 ΔmeeY meeF-FLAG::MLS* | pMUTIN-FLAG | This work |

| | | | |
|---|---|---|---|
| HBAS949 | *trpC2 ΔmeeF meeY-FLAG::MLS* | pMUTIN-FLAG | This work |
| HBAS930 | *trpC2 ΔmneP ΔmneS meeF-FLAG::MLS* | pMUTIN-FLAG | This work |
| HBAS946 | *trpC2 ΔmneP ΔmneS meeY-FLAG::MLS* | pMUTIN-FLAG | This work |

| **Primer name** | **Sequence** | **Purpose** |
|---|---|---|
| meeFcheckF | GTGTCATCCATACGGGTGACAA | Deletion check |
| meeFcheckR | GATTTGAGTGCTTCGCAATCAGCT | |
| meeYcheckF | GGAGGAAGGCCGTTTCTTGA | Deletion check |
| meeYcheckR | CGGATGAAACCGTCTTTGCG | |
| aprEcheckF | CGAGTCTCTACGGAAATAGCGAG | Deletion check |
| aprEcheckR | CTTGTGAAGATTTTCAGAGGCAGC | |
| nprBcheckF | GCAGCTCATACCTCCGCTTATAC | Deletion check |
| nprBcheckR | GATTGGAGCAATCAAACCGTCG | |
| nprEcheckF | CTCTTGAATGAACCACCACATGAC | Deletion check |
| nprEcheckR | ATAGCCACGTGACCTGTAGC | |
| mprcheckF | GCTTGACACTAAAGGAGGGAGATG | Deletion check |
| mprcheckR | GAAGAATGCGTCTGCGAACTG | |
| eprcheckF | CACCCGAGTGAATGTGCTCAT | Deletion check |
| eprcheckR | CATCATTGGGTCTTGCCTGC | |
| bprcheckF | CAGCGATGTTCTGACAAACCATTC | Deletion check |
| bprcheckR | TGCCGTGAGCAAAAGCAAAAG | |
| ftsHcheckF | CTGAGCGCTATCGCAATCTG | Deletion check |
| ftsHcheckR | TACACGATCAGCGGCTCA | |
| ltaScheckF | GCGAAACGTTGATTCGACGG | Deletion check |
| ltaScheckR | GCTGAGGAATTGAGGGCTG | |
| ftsH700F | ATTGTGACGGACGCAAGCGGTGAAATTATC | PCR insertion |
| ftsH700R | TCCTGTCGCAATGACCTTTGGTTCCTGT | |
| aprEFLAGHindIIIF | ATATAAGCTTTCTCACGGCACACATGTAGCCG | Cloning in pMUTIN-FLAG |
| aprEFLAGKpnIR | ATATGGTACCTTGTGCAGCTGCTTGTACGTTG | |
| meeFXbaIF | CAGTTCTAGAAAAGGAGGAAGGATCATTGGACTTTTTACATCATAT TTTGTCTACG | Cloning in pPL82 |
| meeFBglIIR | CGTTAGATCTTTATTCTTCTTTTGAAGCGGCTGTTT | |
| meeYXabIF | ATATTCTAGACAAATCACTGCGCTGCGCATATTATTG | Cloning in pPL82 |
| meeYBglIIR | CGTTAGATCTTTACGCCCGTTCACGGGTGCTG | |
| lmo0991XbaIF | CAGTTCTAGAAAAGGAGGAAGGATCAATGGATACAGCAATGATTTT AGAGTAC | Cloning in pPL82 |
| lmo0991BglIIR | CGTTAGATCTTTATTTAGTTGTTTCTGTTTTTTTCTTAC | |
| lmo0992XbaIF | CAGTTCTAGAAAAGGAGGAAGGATCAATGGATGTTTCTATTTGGG GCGAATATG | Cloning in pPL82[5] |
| lmo0992BglIIR | CGTTAGATCTTCAAACTCTTTTGGTTTTCTTCTC | |

| | | |
|---|---|---|
| BanTerCXbalF | CAGTTCTAGAAAAGGAGGAAGGATCAATGAGTATTTTGCAAGGAATCCTTGATAC | Cloning in pPL82 |
| BanTerCBglIIR | CGTTAGATCTTTATTTATGGTTATTTTTAGTAGCTGCAAC | |
| ftsHXbalF | CAGTTCTAGAAAAGGAGGAAGGATCAATGAATCGGGTCTTCCGTAATACCATTTTT | Cloning in pPL82 |
| ftsHBglIIR | CGTTAGATCTTTACTCTTTCGTATCGTCTTTCTTTTCTTCTGTT | |
| meeF-FLAG-HindIII-F | ATATAAGCTTTGGTGGATCAAGGTGCTTGGCGCGCTTTACCTGGCTTG | Cloning in pMUTIN-FLAG |
| meeF-FLAG-KpnI-R | ATATGGTACCTTCTTCTTTTGAAGCGGCTGTTTGTTCGCGCAC | Cloning in pMUTIN-FLAG |
| meeY-FLAG-KpnI-F | ATATGGTACCAAGAAGACACACATAAAGAGACGAAGCAAAG | Cloning in pMUTIN-FLAG |
| meeY-FLAG-KpnI-R | ATATGGTACCCGCCCGTTCACGGGTGCTGTTTTTTTGTTC | Cloning in pMUTIN-FLAG |
| pMUTIN4-FLAG check-F | ACATCCAGAACAACCTCTGCTAAAATTC | Cloning check; pMUTIN-FLAG |
| ltaS-F-qPCR | TTCAGTTTTCGTAAACAAAGCGC | qRT-PCR |
| ltaS-R-qPCR | ATGCCTTCCGCCTTGTACATTTC | qRT-PCR |
| ltaSa-F-qPCR | CAGCCATTATTATGCTGATTATCG | qRT-PCR |
| ltaSa-R-qPCR | GCCATGATACTGAAAATCCCGTC | qRT-PCR |
| yvgJ-F-qPCR | GCCAATTTGGTTCTGACTGTTATC | qRT-PCR |
| yvgJ-R-qPCR | CTTTTACACTGCTGCCCATATCGCTC | qRT-PCR |
| yqgS-F-qPCR | ATGCTGATTGCCATTTTATTGATGTG | qRT-PCR |
| yqgS-R-qPCR | TGCCAGCAATACAAACGTGAC | qRT-PCR |

| **Plasmid** | **Properties** | **Reference** |
|---|---|---|
| pPL82 | IPTG-induced overexpression construction | [6] |
| pKTH10 | overexpression of AmyQ | [7] |
| pMUTIN-FLAG | tagged genes with FLAG sequence at C-terminal | [5] |
| pBS3C*lux* | luciferase promoter construction | [8] |

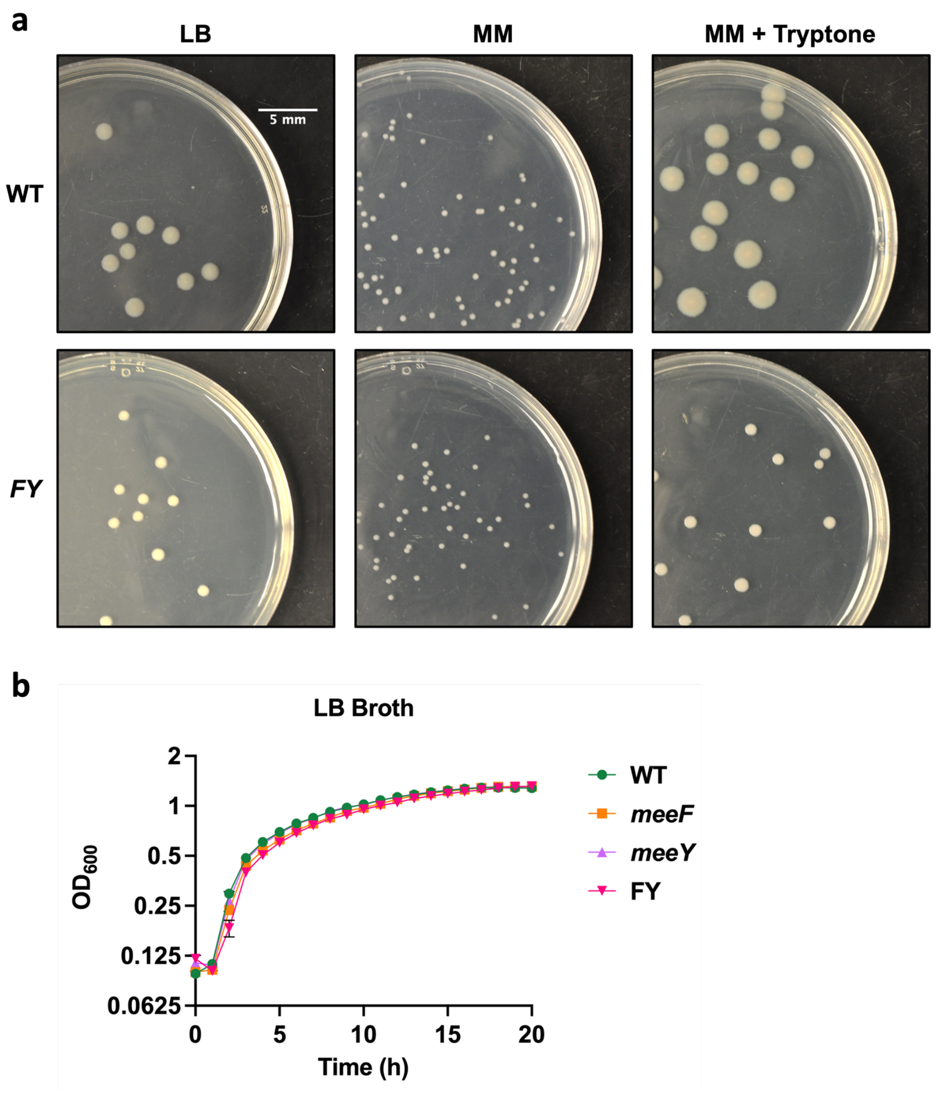


**Figure S1. Comparative growth of WT and FY mutants.** (**a**) Growth of WT and FY mutants on LB and defined (MM) agar plates. Representative pictures of colony size on different medium plates are shown. Scale bar (5 mm) is indicated and applies across all images. (**b**) Aerobic growth in liquid LB medium. Representative growth curves of different strains (WT, *meeF, meeY, FY*) in liquid LB broth was performed. Cultures were incubated at 37˚C and changes in $OD_{600}$ nm was measured every hour in a 96-well plate reader (Bio-Tek).

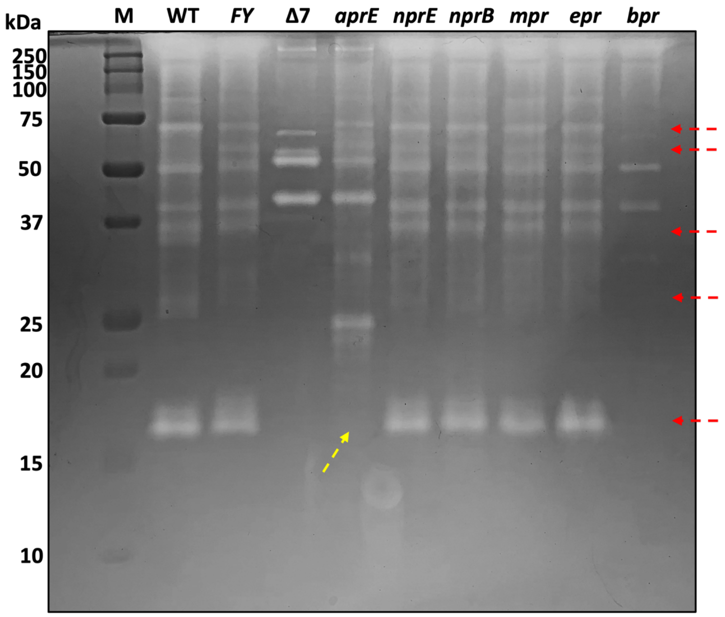


**Figure S2. Extracellular protease activities in the supernatants of protease mutants.** Extracellular protease activities in the supernatants were detected by gelatin zymography. Supernatants were collected from overnight cultures with the same cell number. Higher protease activities correspond to clearer bands on the gel matrix. Note that many of the bands lacking in the *bpr* null mutant were previously proposed to be processed products of the large Bpr protease (red arrows)[9]. The 17 kDa processed Bpr product is also missing in the *aprE* mutant (yellow arrow), suggesting that the AprE protease is involved in Bpr processing, as suggested[9]. The presence of the 17 kDa band in the FY mutant suggests that both Bpr and AprE are still secreted in this strain, although overall protease activity is reduced.

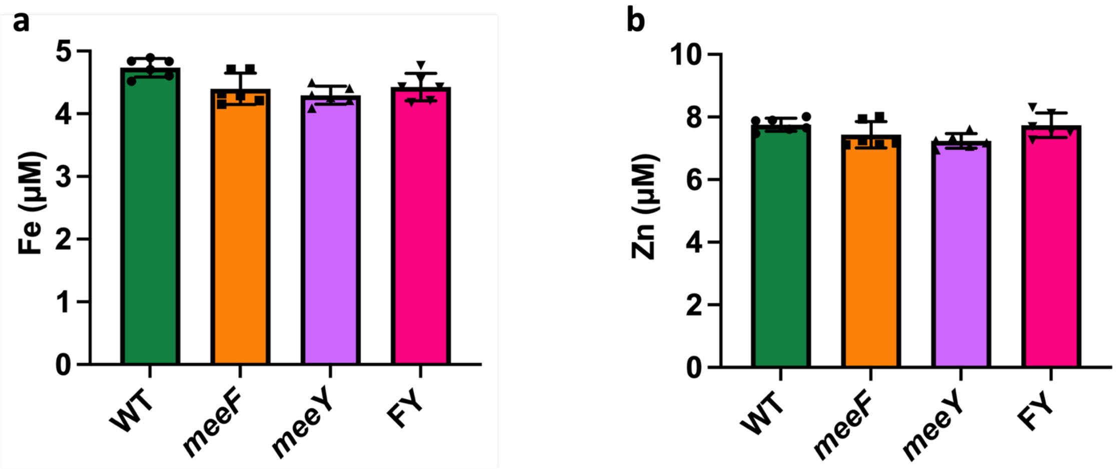


**C**. Table. Metal levels (µM) in LB or spent supernatants detected by ICP-MS

| | LB broth | WT | *meeF* | *meeY* | *FY* |
|---|---|---|---|---|---|
| **Mn** | 0.126±0.009 | 0.022±0.002 | 0.017±0.005 | 0.021±0.003 | 0.0032±0.0009 |
| **Fe** | 5.05±0.62 | 4.73±0.15 | 4.40±0.25 | 4.30±0.14 | 4.43±0.22 |
| **Zn** | 8.20±0.87 | 7.75±0.21 | 7.43±0.42 | 7.24±0.23 | 7.74±0.39 |

**Figure S3. Metal ion levels (µM) measured in cell supernatant fractions.** Overnight grown cells were used to collect supernatants and Fe (**a**) and Zn (**b**) levels were detected using ICP-MS analysis. Data is from three independent experiments and presented as mean ± SD. In contrast with Mn levels (Fig. 2b), there was little change in residual Fe and Zn in the supernatant fractions. (**c**) Summary Table: Samples were collected and analyzed as in Fig. 2b, S3a and S3b. Samples were from three independent experiments and data is presented as mean ± SD.

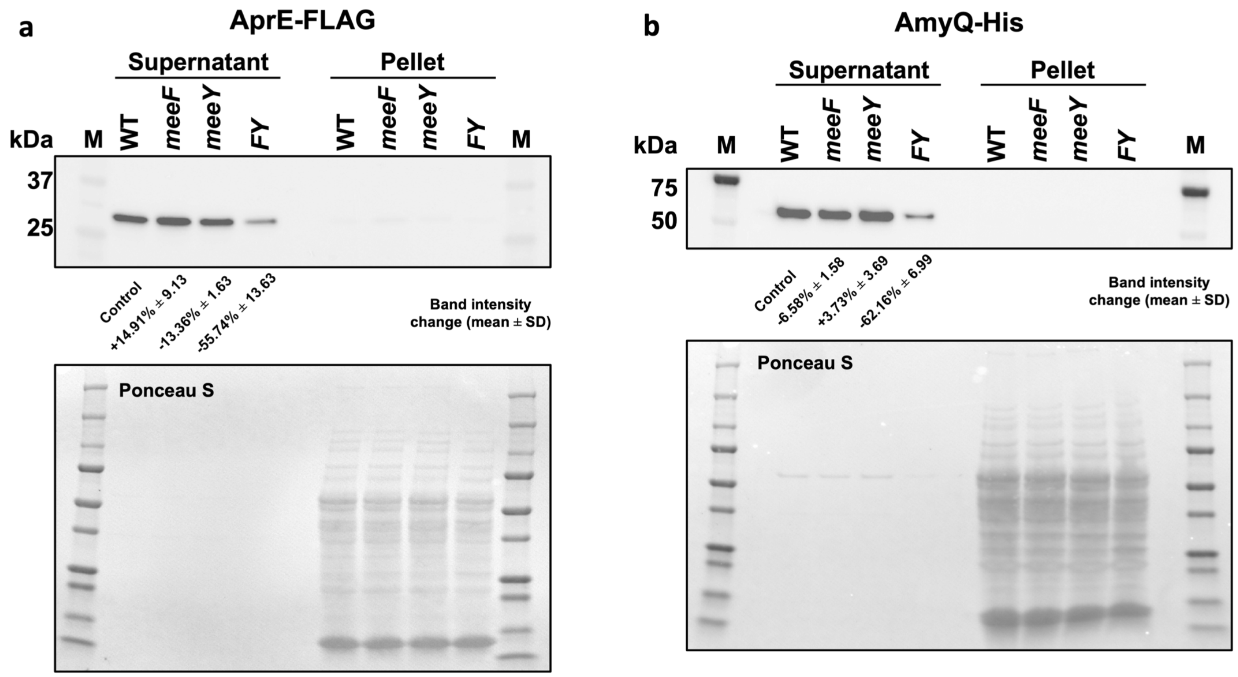


**Figure S4. Effect of *meeF*, *meeY* and FY mutations on AprE and AmyQ secretion**. (**a**) The level of AprE-FLAG was measured using immunoblotting. (**b**) Defective secretion of heterologous (AmyQ-His) protein is depicted for FY. Sample collection was described in Fig 3. The protein membranes were stained by Ponceau after protein transfer. Ponceau-stained images as the loading control were taken using GelDoc gel imaging system. Band intensities were measured from three independent experiments. Band intensity change was calculated as "change = (sample - control) / control * 100%".

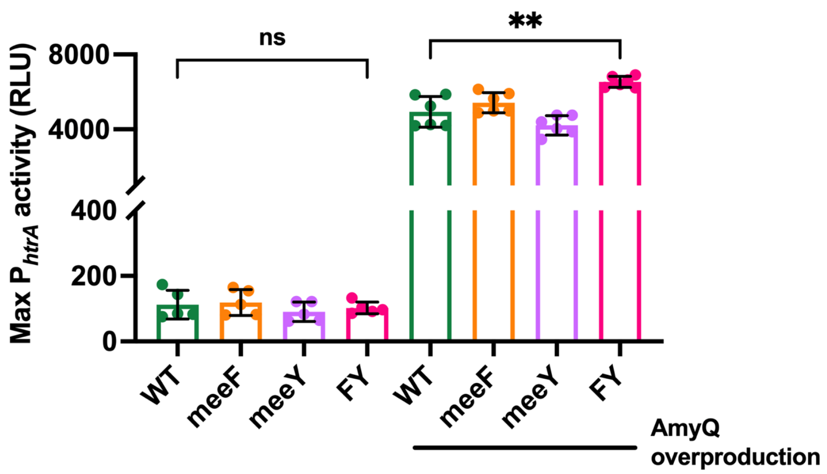


**Figure S5. Maximum P$_{htrA}$-lux promoter activity in stains with or without AmyQ overexpression.** Expression of the CssRS secretion stress response was monitored using an P$_{htrA\text{-}lux}$ transcriptional reporter. Cells were grown in LB medium at 37˚C. AmyQ was overexpressed in the strains from pKTH10[7]. Data are presented as mean ± SD. ns, no significant difference, P=0.6540; **, P=0.0035, P value was calculated using Welch's t test, two-tailed.

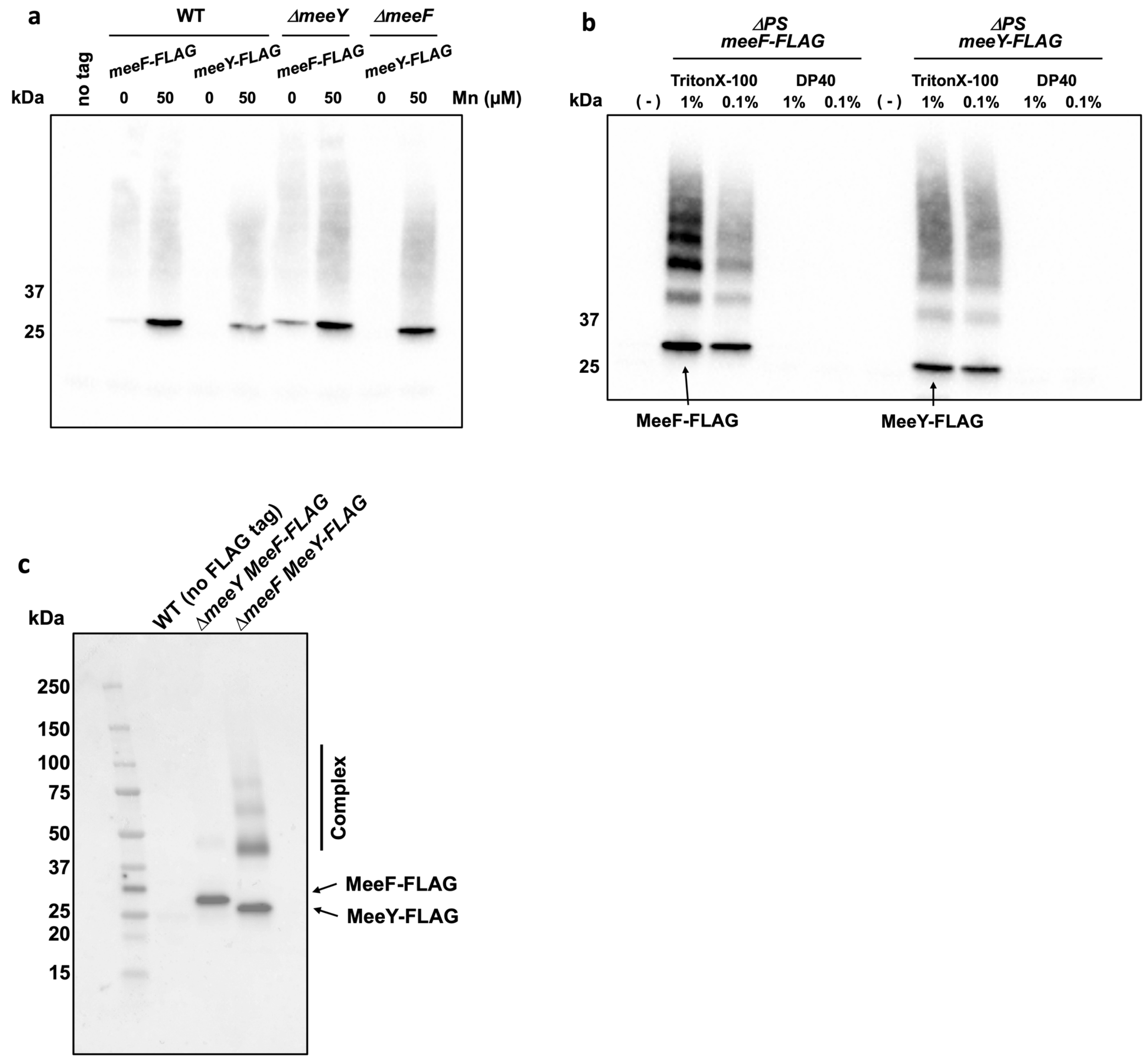


**Figure S6. MeeF-FLAG and MeeY-FLAG in different mutants.** (**a**) MeeF-FLAG or MeeY-FLAG levels in different stain backgrounds as measured using immunoblotting. Cells were collected from overnight cultures with or without 50 μM Mn. (**b**) Representative immunoblot analysis of MeeF-FLAG and MeeY-FLAG co-IP samples in an efflux deficient *mneP mneS* (Δ*PS*) background. Different detergents were used for cell lysis (1% or 0.1% Triton X-100, and 1% or 0.1% DP40). Proteins were eluted from magnetic beads by heating at 95˚C for 10 min. (-) WT is the control without FLAG tag. (**c**) Co-Immunoprecipitation of MeeF and MeeY interact.

Representative Western-blot analysis of MeeF / MeeY - FLAG (bait) is shown following pull down of cell lysates prepared with 1% Triton X-100 using anti-FLAG-magnetic beads and subsequent elution. The WT strain without FLAG tag was used as a control to diminish false positive protein interactions. Proteins (prey) that identified in the immunoprecipitate (and not present with the control, untagged strain) are considered interacting protein partners of MeeF/MeeY-FLAG. The detection of lower mobility bands represents a possible complexes stable during electrophoresis.

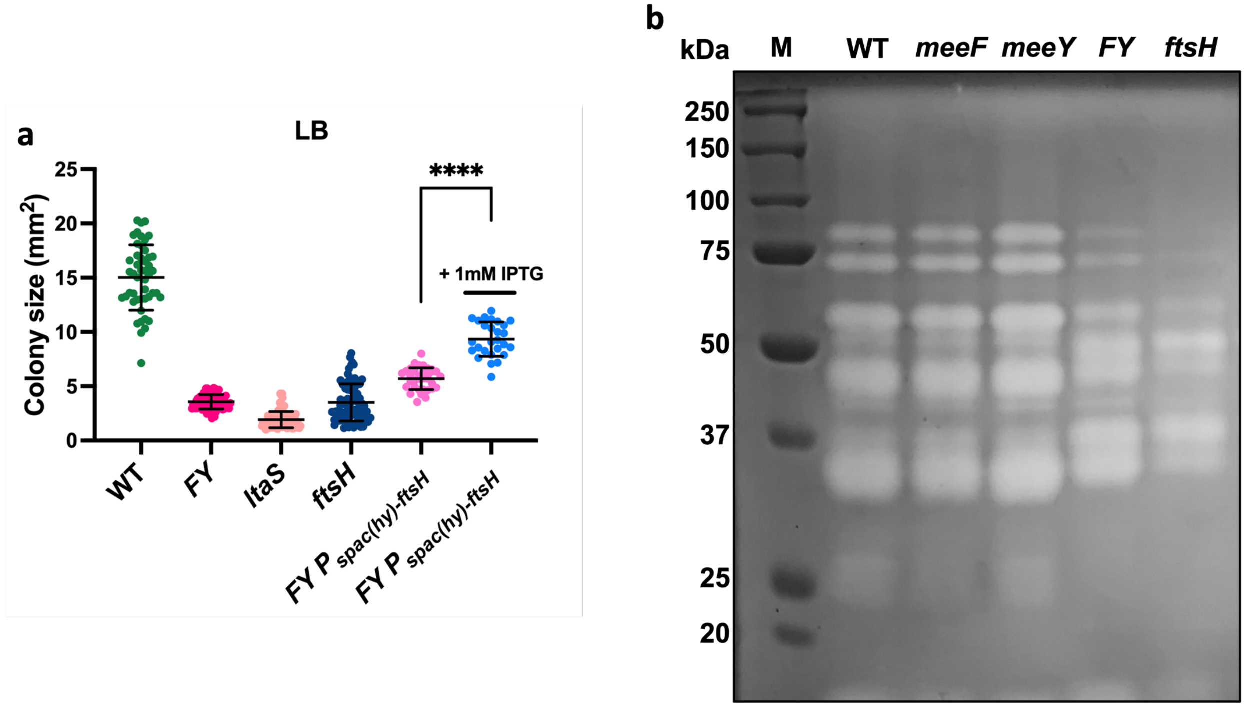


**Figure S7. Comparative phenotypes of mutations affecting protein secretion.** (**a**) Colony size (mean ± SD) of the indicated strains on LB medium. At least 25 isolated colonies from two independent experiments were measured for each strain. Growth of the FY mutant strain was significantly improved upon addition of 1 mM IPTG (**** = P<0.0001). P value was calculated using Welch's t test, two-tailed. (**b**) Protease secretion monitored by zymography. Supernatants were collected from overnight cultures with equal cell density. Higher protease activities represent clearer bands on gel matrix. *ftsH* mutant showed defective protease activities but different pattern compared to FY. M, marker.

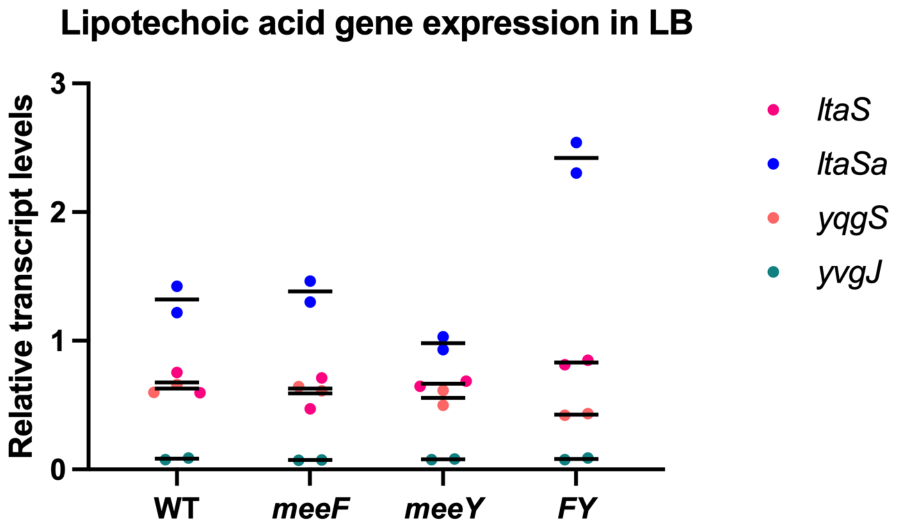


**Figure S8. Fold change of transcription level of LTA synthase genes in different strains.** Transcription levels (mean ± SD) of four LTA synthases or primase *ltaS*, *ltaSa*, *yqgS* and *yvgJ* in different stains (WT, *meeF*, *meeY* and FY). Samples are from two independent experiments. FY mutants showed similar *ltaS* expression and higher *ltaSa* transcription levels compared to WT, which is consistent with the hypothesis that FY mutant has normal expression of *ltaS* but decreased LtaS activity.

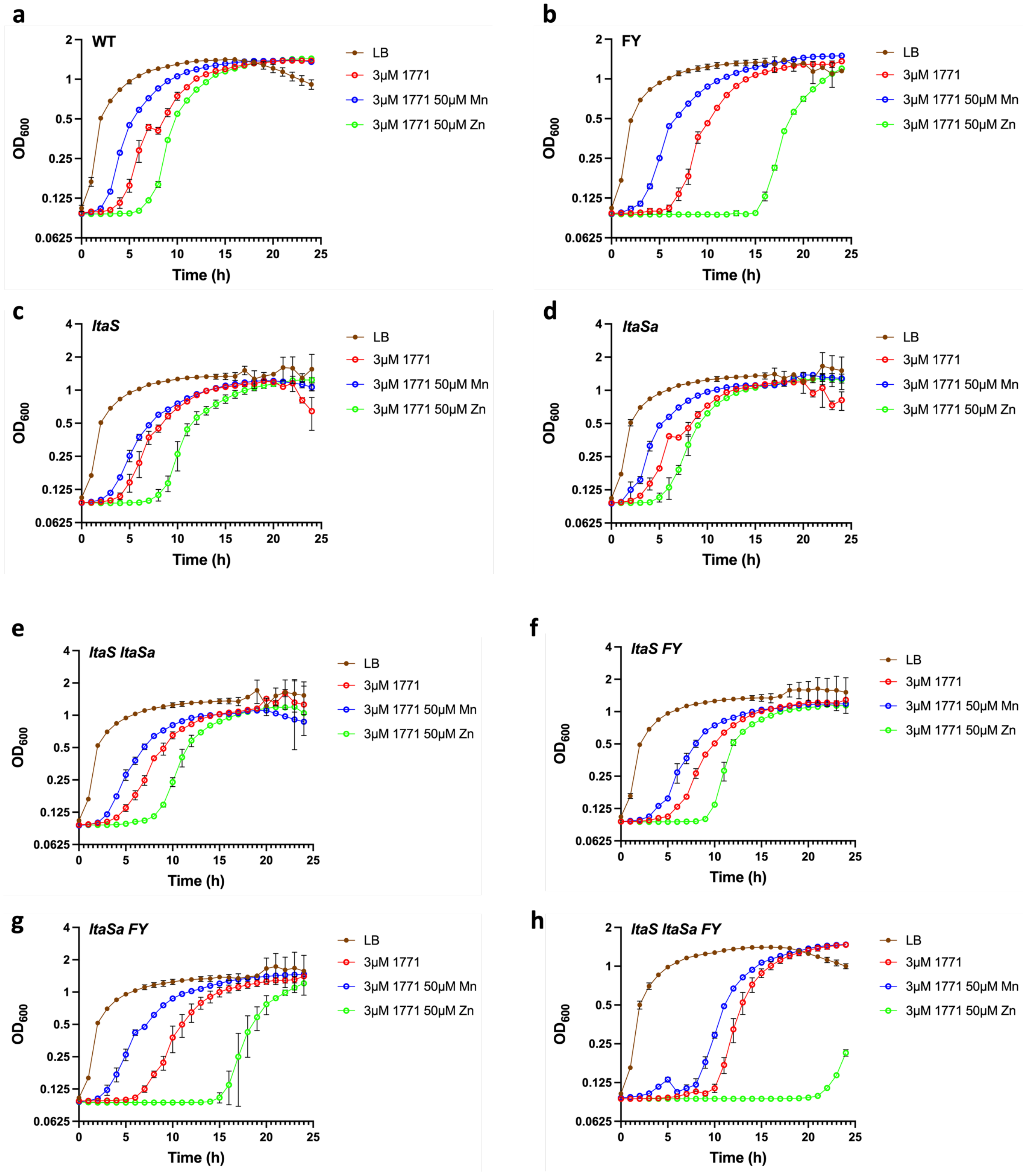


**Figure S9. Effects of metal ions of sensitivity to the LtaS inhibitor 1771.**

(**a**)-(**h**), Aerobic growth of different strains in LB broth with or without 3 μM 1771, 50 μM Mn or 50 μM Zn. Data is representative of three independent cultures and presented as mean ± SD.
